# A QuantCrit investigation of society’s educational debts due to racism, sexism, and classism in biology student learning

**DOI:** 10.1101/2022.05.05.490808

**Authors:** Jayson Nissen, Ben Van Dusen, Sayali Kukday

## Abstract

We investigated the intersectional relationships between racism, sexism, and classism in inequities in student conceptual knowledge in introductory biology courses using a quantitative critical framework. Using Bayesian hierarchical linear models, we examined students’ conceptual knowledge as measured by the Introductory Molecular and Cell Biology Assessment. The data came from the LASSO database and included 6,547 students from 87 introductory courses at 11 institutions. The model indicated that students with marginalized identities by race, gender, and class tended to start with lower scores than continuing-generation, White men. We conceptualized these differences as educational debts society owed these students due to racism, sexism, and classism. Instruction added to these educational debts for most marginalized groups, with the largest increases for students with multiple marginalized identities. After instruction, society owed Black and Hispanic, first-generation women an educational debt equal to 60-80% of the average learning in the courses. These courses almost all (85/87) used collaborative learning and half (45/87) supported instruction with learning assistants. While research shows collaborative learning better serves students than lecture-based instruction, these results indicate it does not repay educational debts due to racism, sexism, and classism.

## Introduction

Science, Technology, Engineering, and Mathematics (STEM) fields have historically marginalized students belonging to groups including persons excluded due to ethnicity or race, women, and first-generation students. This marginalization has resulted in negative outcomes on metrics that are important for student success in STEM fields including test performance, grades, graduation rates, retention in major, course failure, science identity, self-efficacy, and sense of belonging (Asai, 2020, Theobald et al., 2020; Eddy et al., 2014; Ballen et al., 2017; Seymour and Hunter, 2019; NCSES, 2021). Students experiencing these negative outcomes leave at different points along their STEM education progression (Seymour and Hunter, 2019). These losses deny the students fair opportunities to pursue their passions and curiosity. They further act to exclude their communities from having voices and leaders within the scientific communities tasked with addressing many issues society faces. Losing talent and diversity of perspective from this attrition has led education researchers to recognize the existence of the problem, investigate underlying mechanisms, and propose approaches that might help address the problem. In this paper, we operationalize the writings of Ladson-Billings (2006) to describe differences in student performance on the Introductory Molecular and Cell Biology Assessment (IMCA; Shi et al., 2010) in terms of society’s accrued educational debt owed to marginalized students due to racism, sexism, and classism.

## Definitions

To support readers’ interpretation of our research, Table 1 includes a selection of terms and definitions for statistical modeling and equity-related terms that we use.

**Table 1.** Definition of Some Statistical Modeling and Equity-Related Terms We Use in the Manuscript

| Term | Definition |
| --- | --- |
| Hierarchical Linear Model | A linear regression model that accounts for the nested nature of data (e.g., students within courses). Also known as a multilevel model. |
| Bayesian | A statistical system based on probability, rather than frequency. |
| Multiple Imputation | A principled approach to handling missing data that generates multiple complete data sets based on collected data and combines the results to account for the increased variance. |
| Uncertainty (Standard Error) | In Bayesian statistics, the uncertainty provides a probability that a true value falls within a range. The greater that range for a given probability the greater the uncertainty. |
| Equality | When things are equal between groups of people. This can be applied to different measures, such as having equal resources, opportunities, or outcomes. In this work, we focus on equal outcomes. |
| Equity | When a course allocates resources and opportunities according to each person's circumstances to reach equal outcomes. |
| Educational Debt | "Education debt is the foregone schooling resources that we could have (should have) been investing in (primarily) low-income kids, which deficit leads to a variety of social problems (e.g., crime, low productivity, low wages, low labor force participation) that require on-going public investment".<sup>a</sup> This reframes inequities in group outcomes from deficits in student abilities to debts that society owes marginalized students due to racism, sexism and classism. (Ladson-Billings, 2006; 2007) |
| Intersectionality | "Intersectionality means the examination of race, sex, class, national origin, and sexual orientation and how their combinations play out in various settings." (Delgado & Stefancic, 2017) |
| Critical Race Theory | "[Critical race theory] begins with the notion that racism is 'normal, not aberrant, in American society' (Delgado, 1995, p xiv), and, because it is so enmeshed in the fabric of our social order, it appears both normal and natural to people in this culture." (Ladson-Billings, 1998) |
| QuantCrit | A theoretical framework that applies critical race theory to quantitative research. |
| First-Generation | A student whose parent(s) or guardian(s) do not have a bachelor's degree. |

## Literature Review

Looking at the proportions of undergraduate degrees earned by different groups in the biological sciences provides little evidence of racism advantaging White students or sexism advantaging men. White^1^ students earn 54% of the undergraduate degrees in the biological sciences (NCSES, 2021) and make up 54% of the 18-24-year-olds in the United States (NCES, 2019). Women earn 63% of the degrees compared to making up 50% of the 20-24-year-old population (US Census Bureau, 2021). Though Black (7%) and Hispanic students (14%) earned (NCSES, 2021) a much lower proportion of degrees than their 14% and 24% of the 18-24-year-old population (NCES, 2019). Asian Americans made up 6% of the college age population (NCES, 2019) but earned 14.5% of undergraduate degrees in the biological sciences (NCSES, 2021). Metcalf (2015), however, points out that we should look more broadly and critically at equity in who earns life-science degrees and participates in life-science careers.

Trends shift to illustrate advantages for men and White students in data from careers that build on degrees in the life sciences. The proportion of women entering careers as practicing life scientists was 48% in 2019, a large decrease from the 63% of bachelor’s degree earners (NCSES, 2021). White Americans made up 63% of people entering these careers, a marked increase over the 54% of undergraduate degrees in the biological sciences they earned. The proportion of those entering the workforce that identified as Black (3.6%) and Hispanic (9.8%) was also lower than the proportions of bachelor’s degrees, while the proportion for Asian Americans was higher at 21.4%. Similar trends show up when looking at the proportion of PhDs in the biological sciences awarded to women (53%), White (68%), Asian (11%), Black (4%), and Hispanic (8%) students (NCSES, 2021).

Differential impact of opportunities and resources provided in life-science courses for women and students of color may accumulate and contribute to the increased representation of men and White Americans in the workforce and earning postgraduate degrees. Researchers often describe these differences in outcomes as “achievement gaps” (Haycock, 2001). The term promotes a deficit perspective portraying marginalized students as deficient in some domain (e.g., lack of content knowledge, attitudes, or grit; Gutierrez, 2008; Gutierrez & Dixon-Román, 2010, Shukla et al., 2022). Because of the preponderance of this term in education research literature, we use “achievement gap” in this section to remain consistent with the literature. We, however, reframe the differences in group performance as societal educational debts (Ladson-Billings, 2006; Shukla et al., 2022) in our own work. As discussed in our Conceptual Framework, we use societal educational debts to focus on society and institutions as the historical and ongoing sources of these inequalities (Shukla, 2022).

Performance or achievement measurements to assess student learning often include test score data to compare differences among students based on race, gender, or first-generation status. Eddy et al. (2014) studied gender disparities in achievement and participation of undergraduate students in introductory biology and found that female students with the same GPA and racial/ethnic background as their male counterparts scored 0.2 SDs lower (11 points; 2.8%) on overall exam grades. They inferred a lack of preparation and the experience of stereotype threat as possible explanations for these gender disparities. While stereotype threat research often focuses on the way it can suppress performance, stereotype threat also undermines learning (Taylor and Walton, 2011; Mangels et al., 2012; Rydell et al., 2010). Theobald et al. (2020) investigated the achievement gap in introductory STEM courses for undergraduate students and found that students from groups marginalized in STEM had lower performance on exams and lower course passing rates than their peers based on data from 15 studies that included 9,238 students from 51 introductory STEM classrooms. Their analysis focused on students from under-represented minorities (URM) and low socioeconomic status (SES) backgrounds. These results show a consistent difference in outcomes favoring White students, high-SES students, and men.

Eddy et al. (2015) also found that preferences in how one participates in class differ across social identities. Compared to their counterparts, men prefer leadership roles, whereas women prefer collaborative roles and URM, Asian-American, and international students prefer listening roles. Robertson and Hairston (2022) use an example from an introductory physics course to illustrate how common course structures center students seeking leadership roles. This centering reifies Whiteness within STEM courses by marginalizing those students who seek collaborative or listening roles.

In addition to these differences in learning and classroom participation, Metcalf (2015) points out the large difference between men and women moving into senior and upper-level leadership positions across industry, government, and academia. In 2014, 80 biotechnology companies filed initial public offerings, 20% of these companies did not have women in *any* leadership roles and only six had women CEOs. Within the life-sciences in academia, women comprised 46% of assistant professors, 31% of associate professors, and 23% of full professors.

Other strands of research have focused on successful women and Black, Hispanic, Latino, and Latina students in STEM broadly (McPherson, 2017; Whitten, Foster, and Duncombe, 2011; McGee and Bentley, 2017; Ong, 2005; Ong, Wright, Espinosa, and Orfield, 2011). Stanton et al. (2022) explored the strengths Black biology majors drew on to navigate the racial climate at a primarily White institution. They found that Black students experienced a sense of isolation, often faced racist microagressions, and faced covert racist interactions that often occurred off campus with other students. The microaggressions included assumptions that they were poor or first-generation college students, commenting on and touching their hair, and difficulty finding partners for class activities. Many students developed the ability to educate people who said or did racist things as an empowering form of resistant capital that allowed them to challenge the status quo. They also found and created a wide array of spaces to foster their community. These include formal and informal spaces, in person and virtual spaces, and spaces in and outside of their science disciplines. These results show that Black students drew on several forms of cultural capital to support each other and resist exclusion and racism they faced in their education.

Class, or SES, represents differential access (realized or potential) to desired resources or opportunities (Oakes and Rossi, 2003; Cowan et al., 2012). Oakes and Rossi argue researchers can measure class across three domains: material capital, human capital, and social capital. These align with the “big-3” measures of SES common in education research (Cowan et al., 2012): family income, parental educational attainment, and parental occupation.

Talbot (2021) searched the CBE Life Science Education journal for the terms socioeconomic status, socioeconomic, and SES with follow-up searches for Pell, and income. The search returned 53 articles; thirteen of these articles used some measure of SES. Talbot (2021) found that none of these 53 articles defined SES or used a theory based approach to measuring SES (e.g., Oakes and Rossi, 2003; Cowan et al., 2012). The 13 articles that measured SES used Pell eligibility, college debt, first-generation status, or an institutional determination based on multiple factors. Only one study (Wright et al., 2016) detailed the overlap between SES and race; they found that Black, Hispanic, Native American, and Pacific Islander students were much more likely to have a low SES (79-90%) than Asian (19%) and White (3%) students. The studies consistently found achievement differences favoring higher SES students. In some cases, including other factors in their statistical models explained these differences. Wright et al. (2016) found that low-SES students performed similarly to middle and high-SES students on highly structured test questions but performed worse on open-response questions, which they classified as higher on Bloom’s taxonomy. Their analysis controlled for prior achievement. Eddy and Hogan (2014) found differences advantaging White and continuing-generation students in lecture-based courses decreased when the course integrated more structured activities including student-centered activities and graded preparation or review activities. Their analysis controlled for prior achievement. Sellami et al. (2017) found similar results to Wright et al.; adding more course structure in the form of learning assistants reduced differences between under-represented marginalized students and White and Asian students. Sellami and colleagues’ supplemental material show large differences in the raw data across URM, Pell-eligibility, First-generation, and sex in both courses with and without learning assistants. Wilton et al. (2019) found that lower-SES students were retained in their biology major at much lower rates than middle and high-SES students. Low-SES students also had lower grades and a lower sense of belonging.

## Research Questions

Through this research, we aim to apply QuantCrit (Stage, 2007) to reframe the conversation about student learning in biology from a perspective of “achievement gaps” to educational debts society owes to learners from marginalized groups (Ladson-Billings, 2006). This study builds on the existing literature on inequities in biology education by measuring the intersectional impacts of sexism, racism, and classism on biology student learning across multiple institutions. Racism, sexism, and classism are societal systems that divide *actors* into groups and distribute (or produce) power unevenly based on these classifications either through oppression or privilege (Paradies, 2006). The results can support biology programs in examining the extent to which they repay or add to society’s educational debts. To better understand the intersecting roles that sexism, racism, and classism play in shaping biology student learning, we asked the following questions:

1. To what extent have sexism, racism, and classism created educational debts of biology knowledge that society owes students before taking introductory college biology courses?
2. To what extent do introductory college biology courses mitigate, perpetuate, or exacerbate the educational debts that society owes students?
3. How, if at all, does the intersection of sexism, racism, and classism relate to society’s educational debts before instruction and after instruction?

## Conceptual Framework

### Quantitative Critical Race Theory (QuantCrit)

QuantCrit applies the tenets of Critical Race Theory (CRT; Ladson-Billings, 2013; West, 1995; Sleeter & Bernal, 2004; Delgado & Stefancic, 2017), which has focused on qualitative methods, to quantitative research to address social injustices and racial oppression. Scholars in many fields, including education (Ladson-Billings, 1998; Ladson-Billings, 2009) have used CRT to examine oppressive power structures, challenge the ideas of objectivity, and consider the intersectionality of individual’s identities (Ladson-Billings, 2013; Crenshaw, 1990). Below, we describe four principles of QuantCrit (Gillborn, Warmington, & Demack, 2018; López et al., 2018) and the ways we strove to embody them in this investigation:

#### 1. The centrality of oppression

We assumed that racism, sexism and classism are social processes (Byng, 2012) that we must explicitly examine lest our statistical models legitimize existing inequities. We assume educational inequities come from oppressive power structures that cater to students from dominant groups. As such, we follow Ladson-Billings’ (Ladson-Billings, 2006; 2007) framing of inequities in group performance as educational debts that society owes students due to both historical and ongoing marginalization, which we detail below.

#### 2. Categories are neither ‘natural’ nor given

Society shapes the instruments, categories, analytical methods, and interpretations we used and they reflect the hegemonic power structures within society. Our models aggregated students by social identifiers for race, gender, and first-generation college status. These categories do not represent any natural or scientific truth about students, but are social constructs used to maintain hegemonic power structures. The dynamic socially negotiated natures of race, gender, and class do not diminish the genuine effects of racism, sexism and classism associated with them. We reflected this in our writing by naming racism, sexism, and classism as the causes of educational debts identified by the models.

#### 3. Data is not neutral and cannot ‘speak for itself’

Problematic assumptions that obscure inequities and maintain the status quo can shape every stage of collecting, analyzing, and interpreting data (Barr, Gonzalez, & Wanat, 2008). In our analysis, we drew on research and thinking from across fields to use methods that represented the impacts of racism, sexism, and classism, knowing that the data and methods were imperfect. For example, we used AICc to guide our model development (Van Dusen and Nissen, 2022), multiple imputation to handle missing data (Nissen, Donatello, and Van Dusen, 2019), hierarchical models to address the nested nature of the data (Van Dusen and Nissen, 2019), and visualizations of the raw data to support transparency.

#### 4. The importance of intersectionality

The multiple facets (e.g., race, gender) of identity both intersect with each other and with society’s associated power structures to shape students’ experiences and outcomes (Crenshaw, 1990; Harris & Leonardo, 2018, Covarrubias, 2011; Lopez et al., 2018; Rodriguez, Barthelemy, and McCormick, 2022). Collins (2015) refers to intersectionality as a theory that does not have a precise definition that fits within each field or study but instead draws from a set of guiding assumptions. Several of these assumptions motivated this work. Race, class, and gender, are best understood in relational terms because the power relations of racism, sexism, and classism are interrelated. Intersecting systems of power catalyze complex social inequalities. These inequalities in STEM include unequal outcomes, representations, and social experiences that vary across time and STEM domains. These inequalities are unjust.

Our models followed the advice of Schudde (2018) and used interaction terms for the social identifier variables to allow the models to show the relationships between race/racism, gender/sexism, and class/classism. This allowed the models to have an analytical sensitivity to sameness and difference across the intersections of race, gender, and class in introductory biology courses. Using AICc in our model development ensured the models were sensitive to the additional information provided by the interaction effects and prevented our models from being intersectional in name only. We also focused on using the identities students provided, disaggregating student demographics to the extent our data would allow, and avoiding the use of aggregated groups (e.g., URM) that can obscure inequities for both Black and Asian students (Shafer, Mahmood, & Stelzer, 2021).

## Defining Racism, Sexism, and Classism

We adapt Paradies’ (2006) definition of racism to also apply to sexism, classism, and other forms of oppression. Racism, sexism, and classism are societal systems that divide *actors* into groups; actors include people, institutions, laws, etc. Society distributes (or produces) power unevenly based on these classifications either through oppression or privilege. Inferiority and superiority act as inseparable aspects of these oppressive systems. Because these systemic forms of oppression act through asymmetric power imbalances, they only involve the negative differential treatment of those with less power either through their oppression or through the privileging of those with power. This definition excludes the idea of ‘reverse-racism’ harming those from privileged groups. This definition means that differences across groups in conceptual knowledge are racism, sexism, or classism. Increasing or perpetuating these differences favoring White, continuing-generation men from pre to post instruction are the racist, sexist, or classist outcomes of a course.

Racism, sexism, and classism are broad forces with multiple sources. We use them to make explicit that the inequities we measured in this data represent the outcomes of oppressive ideologies and educational structures. Oppression acts like a cable that binds; it consists of numerous strands derived from the entanglement of racism, sexism, classism^2^, and other oppressive ideologies. These strands fall under four domains, often referred to as the four I’s of oppression: three manifestations of oppression (*institutional, interpersonal*, and *internalized*) and the oppressive *ideologies* at their roots (Paradies, 2006). These implicit and explicit ideologies communicate messages about who belongs in biology and where they belong in the field. These four forms of oppression then act on individuals differently based on the intersections of one’s identities with the various social power structures at play within a given context. This work focuses on the intersectionality of identities with the overall power structures within the introductory college biology context which can lead to differential learning outcomes and representation in life sciences professions. While disentangling the four I’s of oppression in introductory college biology courses is beyond the scope of this work, we use this framework in thinking through the implications of our findings and future studies.

### Operationalizing Equity as Societal Educational Debts

We operationalized equity using societal educational debts (Ladson-Billings, 2006) to interpret the findings from an anti-racist perspective that holds racial groups as equals. Figure 1 illustrates our conception of society’s educational debts. Groups of students enter a course with different skill or knowledge distributions due to oppression, such as the systemic underfunding of schools that disproportionately serve poor students or students of color (Meckler, 2019; Ushomirsky & Williams, 2015). Courses can then either mitigate, perpetuate, or exacerbate those educational debts. Complete mitigation fully repays the educational debt to achieve equality of outcomes (Rodriguez et al., 2012; Espinoza, 2007; Lee, 1999). Complete exacerbation, as shown in Figure 1, represents the complete denial of education to the marginalized group. Researchers often present perpetuation of educational debts, a racist maintenance of educational inequalities, as ‘equity of opportunity’, see Rodriguez et al. (2012) for an example and critique. Society’s educational debts are multifaceted and deeply rooted. The full repayment of one educational debt does not imply the repayment of all educational debts to a marginalized group.

**Figure 1.**
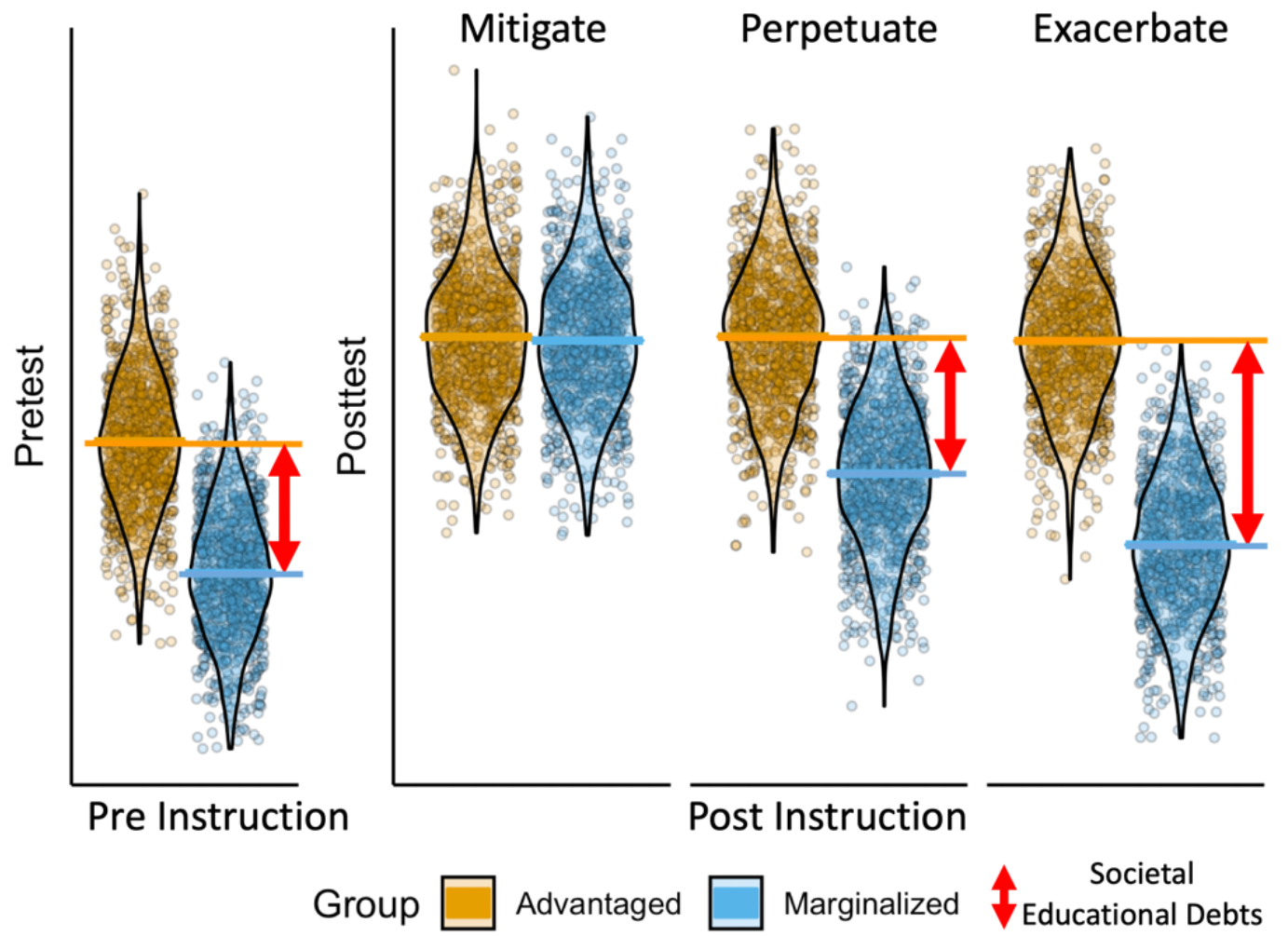
A visual representation of society’s educational debts before and after instruction with three potential outcomes. The figure uses simulated data with the horizontal lines representing mean scores, dots representing individual students, and the violin plot envelopes representing the density of scores. The figure shows how educational debts can be mitigated, perpetuated, or exacerbated. It also shows that in statistical models, educational debts are measures of average differences between groups, not absolute differences between individuals.

Ladson-Billings (2006) introduced the idea of society owing an educational debt to poor, African American, Latino/a, American Indian, and Asian immigrant students similar to the wealth gap between these groups and White men. Like a home, education provides benefits for both those who receive the education and for their children. Thus educational debts accrue both within and across generations. While Ladson-Billings’ 2006 address on educational debts focused on race and class with limited attention to gender, her broader work focuses on the intersections of race, class, and gender (see for example Ladson-Billings, 2009a). And, intersectionality leads us to extend the concept of societal educational debts to racism, classism, and sexism in this work. As Metcalf (2015) points out, the numeric representation of women and underrepresented minorities in biology degrees has not eliminated gender disparities in experiences, opportunities, and persistence within biology education and biology careers. We apply a QuantCrit framework through an educational debts lens represented in Figure 1 to interpret our findings to understand how educational outcomes across the intersections of race, gender, and class mitigate, perpetuate, or exacerbate privilege and oppression in introductory biology courses.

Educational debts in student outcomes, including conceptual knowledge, are power differentials across races, genders, and classes of students. Based on our definition of racism, sexism, and classism, these educational debts are racist, sexist, and classist outcomes of an educational system disproportionately awarding power to men, White, and continuing-generation students. We explicitly state that these educational debts are caused by and due to racism, sexism, and classism to align with our theoretical framework and the research we have reviewed showing the extent of these issues. In light of the extensive research showing racial, gender, and class inequities in STEM, claiming ‘an association with race’, for example, rather than ‘a result of racism’ acts to obscure the systems of oppression that create them rather than to clarify the limitations of an individual study.

### Positionality

The unique experiences and perspectives of the researchers on this team influenced and informed our work. Our identities span genders, races, and disciplinary expertise. To contextualize the research’s perspectives, we provide positionality statements for each author:

The following is the first author’s positionality statement Identifying as a white, cisgender, heterosexual man provides me with opportunities denied to others in American society. My experience growing up poor and serving in the all-male submarine service motivated me to reflect on and work to dismantle oppressive power structures in science. I brought perspective to this work on identity that was shaped by my having a PhD in physics, doing education research and being a white man.

The following is the second author’s positionality statement: I identify as a White cisgender, heterosexual man. I was raised in a pair of lower-income households, but I now earn an upper-middle-class income. I was a continuing generation college student and hold a bachelor’s degree in physics, master’s degree in education, and a Ph.D. in education. My perspective has been informed by my experiences as a faculty member at a teaching-intensive, Hispanic serving institution where I had the privilege to teach and mentor minoritized students. I am currently a faculty member at a research-intensive, predominantly-White institution where I try to use my position and privilege to dismantle oppressive power structures. As someone who seeks to serve as a co-conspirator, it is easy to overlook my privileges. One way that I try to broaden my perspective by soliciting feedback and developing collaborations with peers with different lived experiences than my own.

The following is the third author’s positionality statement: I identify as an Indian cisgender, heterosexual woman. I grew up in a middle-class income household. I was a continuing-generation college student and hold a Bachelor’s degree in Biology and Life Sciences, a Master’s degree in Biochemistry, and a Ph.D. in Biochemistry and Molecular Biology. I am currently a teaching faculty member in a STEM department at a research-intensive Historically White Institution. I hold some privileged and some marginalized identities which inform my teaching, research, and service endeavors. Through my work, I seek to understand the nature of oppression, amplify voices of the oppressed, and advance equity through disrupting oppressive systems.

## Methods

### Data Collection, Cleaning, and Imputation

Student and course data used in this study came from the Learning About STEM Student Outcomes (LASSO) platform research database (Van Dusen, 2018). The LASSO platform automates the administration, scoring, and analysis of research-based assessments online for educators across the STEM disciplines. Several studies have looked at the reliability of collecting data online using low-stakes, research-based assessments. Using a randomized control trial design, Nissen et al. (2018) found that student scores collected with LASSO were similar to scores collected on paper in class. Nissen and colleagues also found that instructors could achieve similar participation levels if they provided participation credit and in-class and email reminders to complete the assessments. Bonham (2008) used a matched sample of students who completed in class and outside of class assessments. Bonham concluded there was no significant difference between the two types of data. He also found that less than 2% of the students copied text from the assessment or systematically used other applications on their computer. Wilcox and Pollock (2019) also concluded that scores were comparable between online and in class low-stakes assessments.

The LASSO research database includes anonymized student and course data from students who consented to share their data. Most educators administered the IMCA as a pretest during the first week of class and as a posttest during the last week of class. The analyzed data came from 6,547 students in 87 first semester introductory college biology courses at 11 institutions. These institutions included two masters institutions and nine doctoral institutions, four of which were very-high research. Two of the institutions were Hispanic serving. Forty-seven of the courses used the Learning Assistant Model (Otero, 2015; Barrasso and Spilios, 2021) to implement collaborative instruction. Eighty-five of the 87 courses reported using collaborative learning. Because the data used for the research contained no identifiable information, it was exempt from IRB review.

Instructors completed a brief survey when setting up their course in LASSO about the pedagogy in their course that includes details on how the LAs supported the course. Of the 87 courses, 61 provided responses to the most recent version of this survey. Figure 2 shows these responses with 54 of the courses reporting students worked in small groups, worked together, or the use of interactive lecture nearly every class or multiple times per class. Two courses reported using all three of these strategies less than weekly though these two courses did report using collaborative learning.

**Figure 2.**
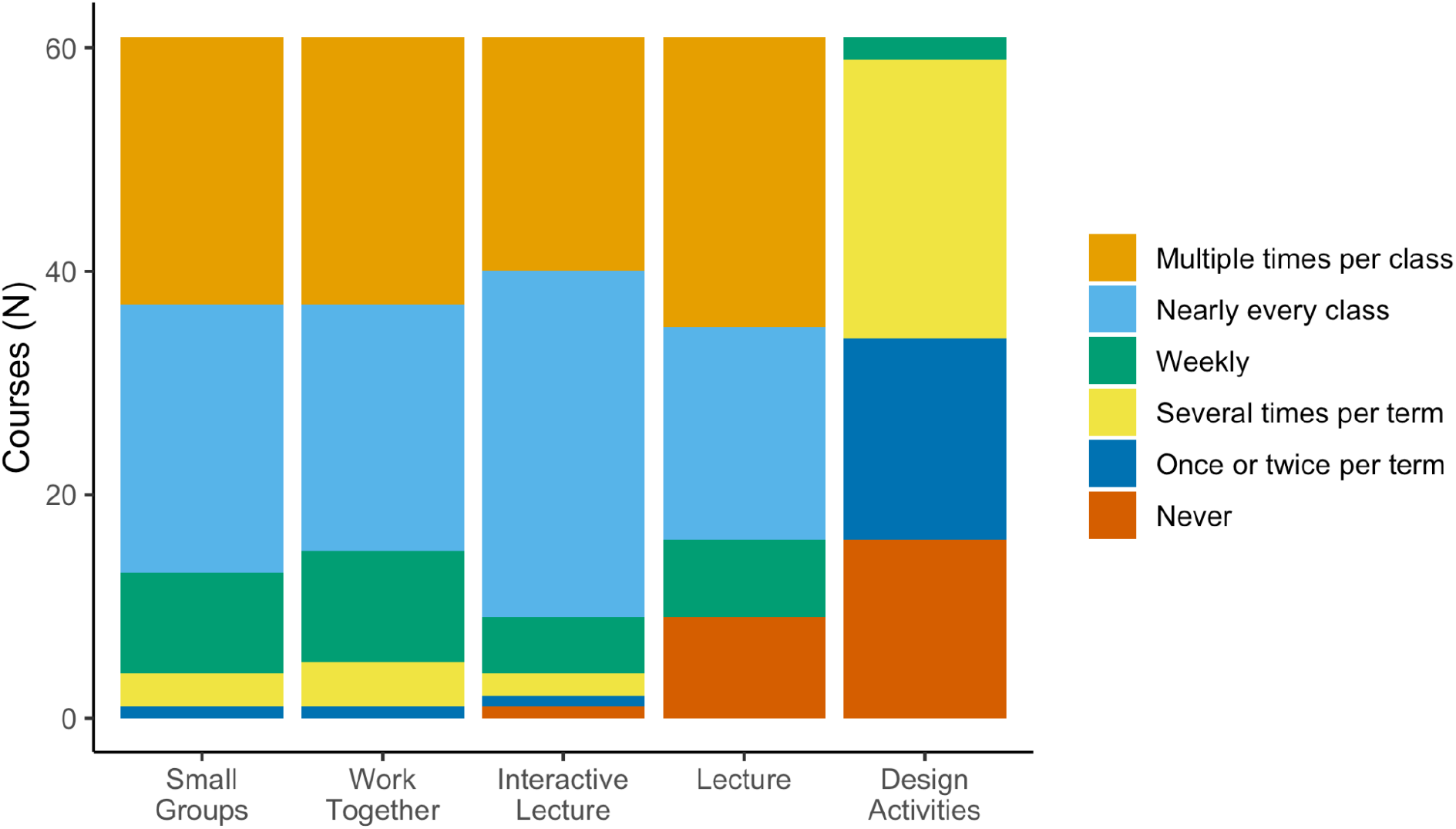
The strategies that instructors reported intending to use in their course for 61 of the 87 courses that completed the most recent version of the pedagogy survey in the LASSO platform. Design activities were activities or experiments that students designed themselves and were uncommon.

The survey also asked instructors about the primary and secondary roles of LAs in their courses, shown in Figure 3, and “How many minutes/week do you plan on meeting with your LAs outside of class, planning for the next week?” The LA model provides structure to support instructors in implementing research-based pedagogies that fit their curricular needs. In the LA model, institutions hire undergraduate students to support instructors using evidence-based, student-centered learning practices in their courses. The LAs take a pedagogy course and pedagogical content knowledge in preparation sessions with the faculty to support them as effective near-peer educators in the classroom. Instructors can implement the LA model across different components of their courses. Instructors primarily planned on LAs supporting their lecture sections or facilitating optional sessions outside of required class activities. Instructors planned on using LAs in mandatory recitations and laboratories much less frequently. Planned meetings with LAs ranged between 20 and 120 minutes outside of class with an average of 70 minutes.

**Figure 3.**
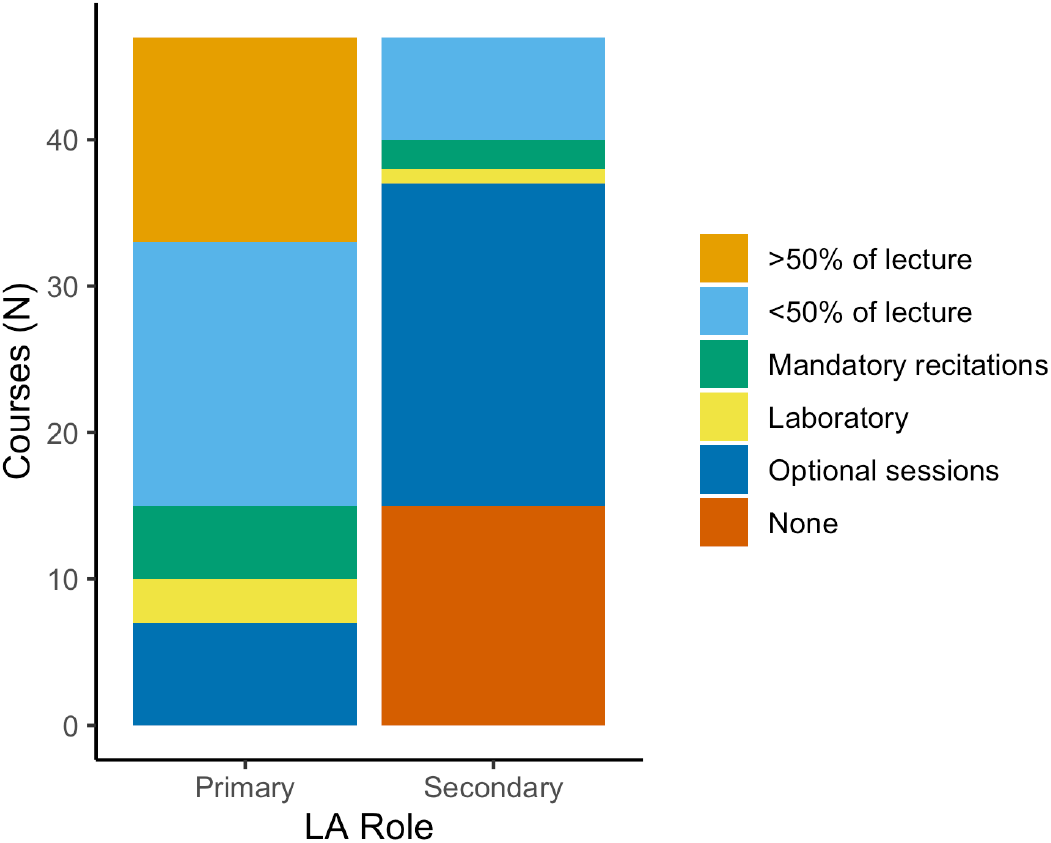
Instructors reported primary and secondary roles for the LAs in their course.

To clean the data, we removed the pretest or posttest score if the student took less than 5 minutes on the assessment or answered less than 80% of the questions. We removed any courses that administered both a pretest and posttest with less than nine pretests and nine posttests and over 60% missing data on either the pretest or posttest. We included courses that only administered the pretest or posttest if they had more than 9 completed tests. After cleaning the data, we used hierarchical multiple imputation (HMI) with the hmi (Speidel, Drechsler & Jolani, 2018) and mice (van Buuren et al., 2015) packages in RStudio V. 1.2.5042 to impute missing data. We imputed values for missing pretest and posttest scores and first generation status. HMI provided a principled method for handling missing data that maximized statistical power and minimized bias while accounting for the hierarchical structure of the data (Allison, 2001; Buhi, Neilands and Goodson, 2008, Manly & Wells, 2015; Nissen, Donatello & Van Dusen, 2019). For the students who provided a pretest, a posttest, or both, 11% were missing the pretest and 33% were missing the posttest. These rates fall within the range of missing data on concept inventories (Nissen, Donatello & Van Dusen, 2019). The LASSO platform added the first-generation college student question during the Fall of 2019. Of the 6,457 students in the dataset, 2,720 (42%) answered the question. To include the variable in the model, we imputed data for students with missing responses. The imputation model included a dependent variable for the posttest and accounted for the pretest score and social identifier variables and nested the students within courses. We included the disaggregated descriptive statistics and plots of student scores in the Supplemental Material with a separate table for the students who provided their first-generation status.

### The Introductory Molecular and Cell Biology Assessment

Shi and colleagues (2010) developed the Introductory Molecular and Cell Biology Assessment (IMCA). They designed it to measure student learning of core concepts in college-level introductory molecular and cell biology courses. Interviews with biology faculty identified topics commonly covered in molecular and cell biology courses. This led to the instrument assessing nine topics: 1) evolution, 2) viruses, bacteria, and eukaryotic cells, 3) structure of macromolecular building blocks, 4) how water affects three-dimensional structures and stability, 5) the impact of thermal energy on reaction rates, 6) solute diffusion and transport, 7) cellular matter and energy flow, 8) storage, replication, and transmission of genetic information, and 9) gene expression.

Shi et al. began item development with student interviews on each topic to identify commonly held beliefs about them. From these interviews, they developed multiple-choice questions with commonly held incorrect ideas as distractors. They then interviewed students about the new items to establish instrument validity. Finally, 25 biology experts reviewed the instrument. Using data from over 1,300 students and three institutions, Shi and colleagues provided evidence for the item and instrument validity.

### Model Building

We developed models of student’s biology conceptual knowledge on the pretest and posttest, described by *score*_*ijk*_ in the final model. The models were 3-level hierarchical linear models (Figure 4) with assessment data (IMCA scores) in the first level (*i*), student data in the second level (*j*), and course data in the third level (*k*). Using hierarchical linear models accounted for the nested nature of the data (Woltman *et al*., 2012; Van Dusen & Nissen, 2019). 5.7% of the variance occurred at the 3rd level (course), 6.7% at the 2nd level (student), and 87.6% at the 1st level (assessment), indicating that a 3-level model was appropriate for modeling the data (Raudenbush & Bryk, 2002). We used Bayesian analysis (Woodworth, 2004) to run the models and pool the results for the imputed datasets using the rstan (Stan Development Team, 2016) and brms (Bürkner, N.D.) packages in R. We discuss our rationale for using Bayesian, rather than frequentist, statistics in the Model Interpretation and Uncertainty section. The model parameters were fit using the penalized least squares method, four chains, 1,000 iteration burn-in, and 2,000 total iterations.

**Figure 4.**
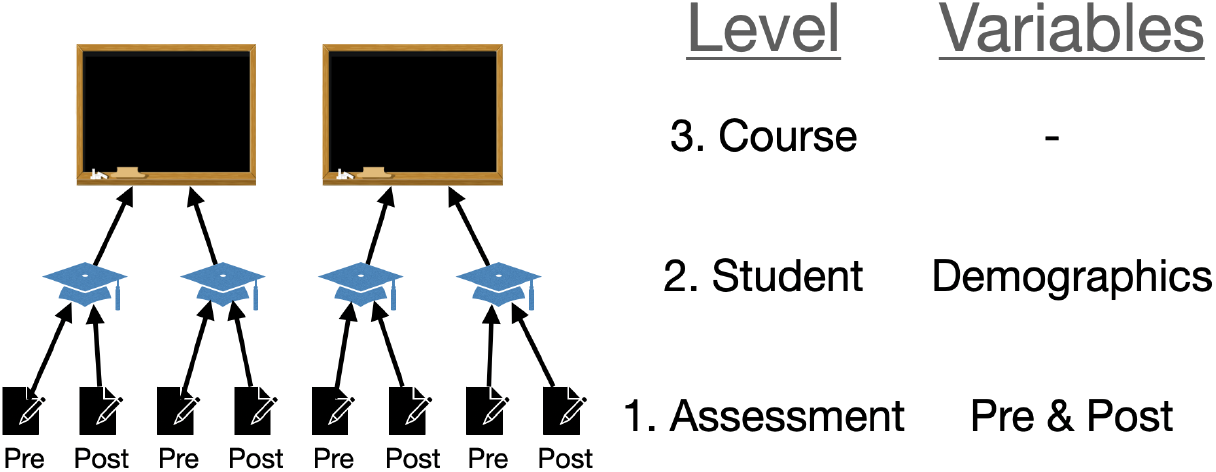
3-level structure of the data and model with the variable categories included in each level.

The data set included social identity data for gender, race, and ethnicity. The social identity questionnaire for the LASSO platform has changed over the period of data collection. We include the most recent questionnaire in the Supplemental Materials. It provided seven gender options, six ethnicity options, 17 race options, and four parental education options. Students could select multiple responses, write in a response, or select prefer not to answer for the gender, ethnicity, and race questions.

Previous social identity surveys on LASSO have conflated gender and sex, which may have shaped how some students responded to these questions. We grouped together students who identified as male or man and those who identified as female or woman. We use the terms men and women to indicate these groups in our model.

To determine what social identity variables to include in the models, we first used a general principle to only investigate scores for populations with at least 20 students total (Simmons, Nelson & Simonsohn, 2011). Following this principle meant that we did not include variables for transgender, Hawaiian or Pacific Islander, or Native American or Alaskan Native in our models. Because removing the students with these identities could have biased the course-level results and because some students did not include a gender or race, we combined these students into two categories: gender other and race other. Our model’s final variables (shown below) included *woman-*_*jk*_, *gender_other*_*jk*_, *Black*_*jk*_, *Asian*_*jk*_, *Hispanic*_*jk*_, *White*_*jk*_, and *race_other*_*jk*_. We included interactions between variables whenever a population had more than 20 students but not for the race other and gender other groups. The model included interaction terms between *Hispanic*_*jk*_ and *White*_*jk*_ and between *woman-*_*jk*_ and each of the model’s racial groups. The first-generation college variable (*FG*_*jk*_) interacted with all the other social identifier variables other than *gender_other*_*jk*_ and *race_other*_*jk*_. Across the intersection of gender, race, and first generation status, two groups fell below our threshold of 20 students: first generation Black men and continuing generation Hispanic men, see Supplemental Table 2. We included the interactions for consistency and will discuss the limitations this introduced in the results.

The models also included several variables not related to social identity. We included the variable *posttest*_*ijk*_ to identify shifts from pretest to posttest and the variable *retake*_*jk*_ to identify whether a student has previously taken the course. Including *retake*_*jk*_ has improved model fit in our prior work (Van Dusen and Nissen, 2022) and did so in this work. We interacted *posttest*_*ijk*_ with all the social identity terms.

We identified our final model using Akaike information criterion corrected (AICc; Johnson & Omland, 2004; Burnham & Anderson, 2004) calculated by the dredge function in the MuMin package (Barton, 2009) in R. We used AICc values because other common techniques for model selection such as coefficient p-values, additional variance explained, and more restrictive information criterion (e.g., Bayesian information criterion) can eliminate social identifier variables or their interactions even when those variables or interactions represent large differences between groups (Van Dusen and Nissen, 2022). Burnham and Anderson (2002) advise against strict cutoffs for model selection. They argue that models within 2 points of the lowest AICc value have equally strong fits and models within 8 AICc points are worth considering. The supplemental material includes the ten models with the lowest AICc scores. For our final model, we only removed the interaction between retake and the pretest, which would be *β*_1(22)*k*_in the final model below, because including the interaction only decreased the AICc by 0.3 points and the interaction was not directly related to our research questions. We left in all other terms in our final model because the AICc scores indicated that removing interaction terms related to our research questions would increase the AICc scores between 3.8 and 9.0 points as shown in the supplemental material.

## Final model

Level-1 equations (assessment-level)

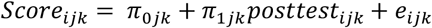

Level-2 equations (Student-level)

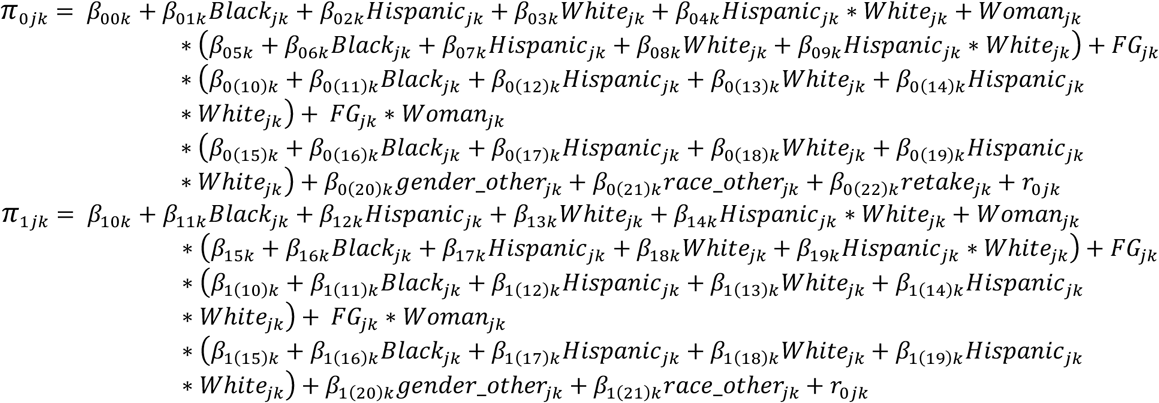

Level-3 equations (Course-level)

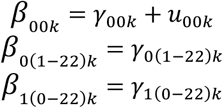

Woltamn (Woltman et al., 2012) provides a detailed description of HLM equations, which we will cover briefly here. The subscripts, for example *Score*_*ijk*,_ refer to the ith assessment in the jth student in the kth course. In the level-1 equation, the π_0*jk*_ term represents the score before instruction. The π_1*jk*_ term represents the shift in scores from before to after instruction. The *e*_*ijk*_ term represents the assessment-level error for a specific score, is the difference between the predicted and actual values, and is analogous to the ε term in standard linear regressions. In the level-2 equation, the *β*_00*k*_ term represents the intercept for scores before instruction. The *β*_10*k*_ represents the intercept for the shift in scores from before to after instruction. The *β*_0(1-22)*k*,1(1-22)*k*_ terms are the coefficients for the respective variable in the model. The *r*_0*jk*_ term represents the student-level error for each student and allows the intercept to vary across each student. In the level-3 equation, the ϒ_00*k*_ term is the intercept for the kth course. The ϒ_0(1-22)*k*,1(0-22)*k*_ terms represent the slopes (e.g., the regression coefficient) for each variable for the kth course. The *u*_00*k*_ term represents the course-level error and allows the intercept to vary across each course. The model is a fixed slope model since the slopes, π_1*jk*_ and *β*_0(1-22)*k*,1(1-12)*k*_equations do not include *r* or *u* variables.

To check the model assumptions, we used visual inspections instead of sensitivity analysis because of its computational difficulty (Gelman & Meng, 1996). Visual inspection showed convergence for all variables.

### Model Interpretation and Uncertainty

To interpret the size of the educational debts, we use 16.3 percentage points as the overall average gain from pre to post-instruction given in the descriptive statistics representing one average term of learning. We reasoned that the average shift over one term provided context for interpreting the educational debts.

We do not use or present p-values in this article. P-values have been misused and misinterpreted throughout the research literature (Ahmrhein et al., 2019; Goodman, 1999). The use of p-values in research has been problematic enough for the American Statistical Association to put out an initial statement on its use in 2016 (Wasserstein & Lazar, 2016) and a second one in 2019 (Wasserstein et al., 2019) that was accompanied by 43 publications offering alternative solutions. Cohen may have put it mostly bluntly in his 1994 article on the misuse of p-values. “After 4 decades of severe criticism, the ritual of null hypothesis significance testing - mechanical dichotomous decisions around a sacred .05 criterion - still persists” (Cohen, 1994, p. 997). In this work we embrace Wasserstein et al.’s (2019) recommendation of accepting uncertainty that goes beyond passing a simple go-no-go test.

To account for uncertainty in the model, we used the standard error for each predicted score to create confidence intervals. We took two steps to prevent these confidence intervals from replicating the shortcomings of p-values (Gigerenzer, 2004; Greenland, 2019; Amrhein et al., 2019). First, we used Bayesian statistics (Woodworth, 2004) rather than frequentist statistics. Measures of uncertainty in frequentist statistics are derived from the probability of a dataset (or a more extreme dataset) given a model (P(data|model)). Unfortunately, this is not a measure of how confident we should be about a model’s estimate; rather it estimates the compatibility of the estimate with the data. Measures of uncertainty in Bayesian statistics, however, are the probability of a model given a dataset (P(model|data)). While researchers often conflate these two probabilities(Gigerenzer, 2004), only the Bayesian measures of uncertainty represent how confident we should be about model estimates. Bayesian models also have the added benefit of their uncertainty measures not depending on large-N approximations, unlike confidence intervals in frequentist analysis (Woodworth, 2004; Krushcke, 2021).

The second way we attempted to prevent the misinterpretation of our findings was by not using error bar overlap as a go-no-go test. If the error bar for two predicted scores did not overlap, we considered those differences robust and likely to occur in future studies. When the error bars overlapped, the differences were less robust and could have been a unique artifact of this data. In this case, we drew on the consistency, or lack thereof, of similar comparisons (e.g., all comparisons of men and women) to inform the robustness of the results.

Figures 6,7,8, and 10 use the data represented in Figure 5 to compare the estimated scores for two groups. In these figures, the error bar represents the standard error from both estimates. Table 2 shows the estimated scores and their standard errors. If this combined uncertainty does not contain zero (i.e. the uncertainties in Figure 5 and Table 2 don’t overlap) then we were very confident the difference is robust. When the error bars included zero, we used the approach discussed in the prior paragraph.

**Table 2.** Predicted pretest and posttest scores from our Bayesian model disaggregated by race, first-generation status, and gender.

| Social Identifiers |  |  | Pretest (%) |  | Posttest (%) |  |
| --- | --- | --- | --- | --- | --- | --- |
| Race | Class | Gender | Score | SE | Score | SE |
| Asian | First-generation | Women | 44.1 | 1.3 | 57.7 | 1.4 |
|  |  | Men | 46.1 | 1.7 | 60.8 | 2.0 |
|  | Continuing-generation | Women | 45.8 | 1.2 | 60.5 | 1.3 |
|  |  | Men | 47.0 | 1.4 | 63.3 | 1.7 |
| Black | First-generation | Women | 41.3 | 1.6 | 52.7 | 1.6 |
|  |  | Men | 43.2 | 3.1 | 55.3 | 3.7 |
|  | Continuing-generation | Women | 41.7 | 1.4 | 55.5 | 1.5 |
|  |  | Men | 45.4 | 2.3 | 58.8 | 2.4 |
| Hispanic | First-generation | Women | 40.3 | 1.6 | 55.1 | 1.7 |
|  |  | Men | 41.4 | 2.1 | 63.1 | 2.3 |
|  | Continuing-generation | Women | 40.5 | 2.5 | 59.9 | 3.5 |
|  |  | Men | 41.2 | 3.2 | 65.3 | 3.1 |
| White | First-generation | Women | 40.8 | 1.1 | 56.1 | 1.1 |
|  |  | Men | 44.0 | 1.6 | 61.8 | 2.0 |
|  | Continuing-generation | Women | 42.1 | 1.0 | 59.3 | 1.0 |
|  |  | Men | 45.7 | 1.0 | 65.4 | 1.1 |
| White Hispanic | First-generation | Women | 39.6 | 1.3 | 55.9 | 1.8 |
|  |  | Men | 43.5 | 2.1 | 61.2 | 2.2 |
|  | Continuing-generation | Women | 41.4 | 1.7 | 59.5 | 1.7 |
|  |  | Men | 41.0 | 2.1 | 63.5 | 2.1 |

**Figure 5.**
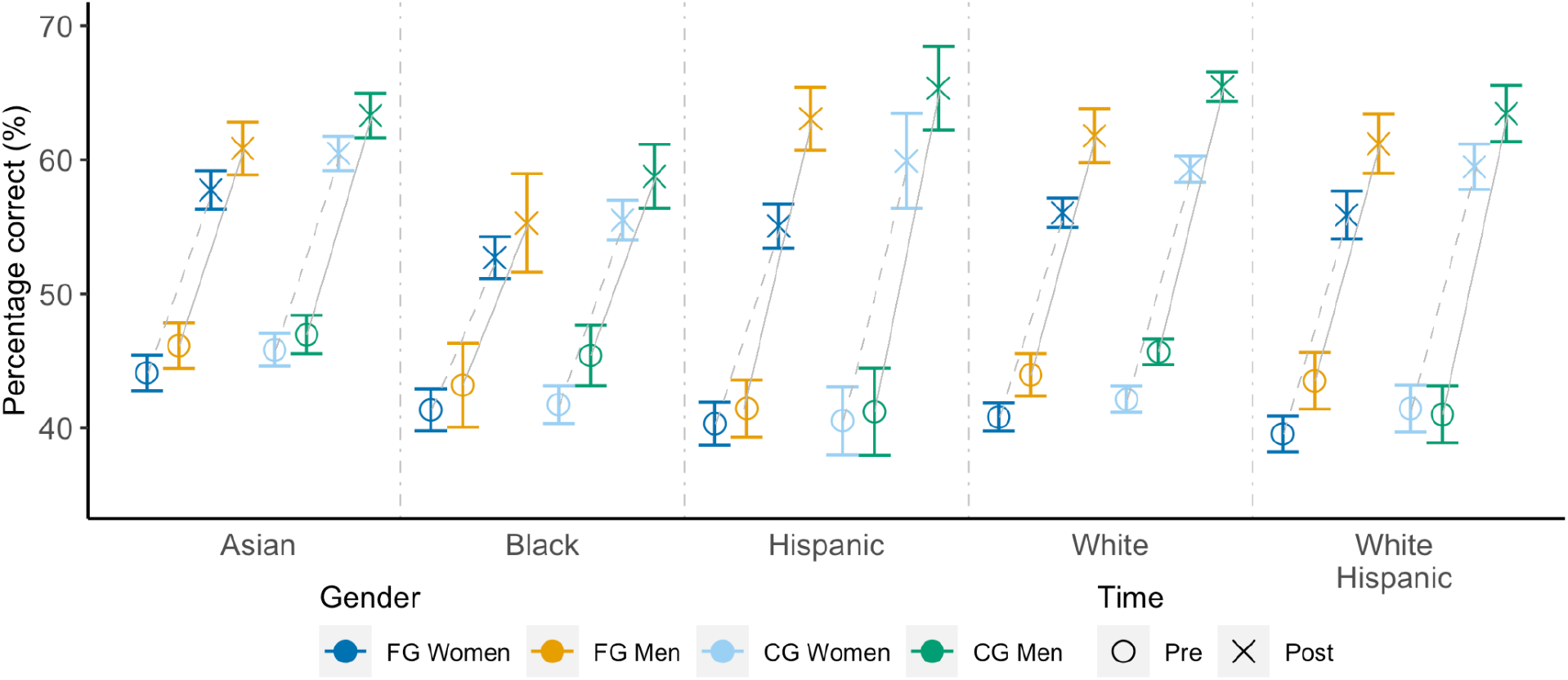
Predicted pretest and posttest scores from our Bayesian model disaggregated by race, gender, and first-generation status (first-generation [FG] and continuing-generation [CG]). Error bars are +/- 1 S.E.

**Figure 6.**
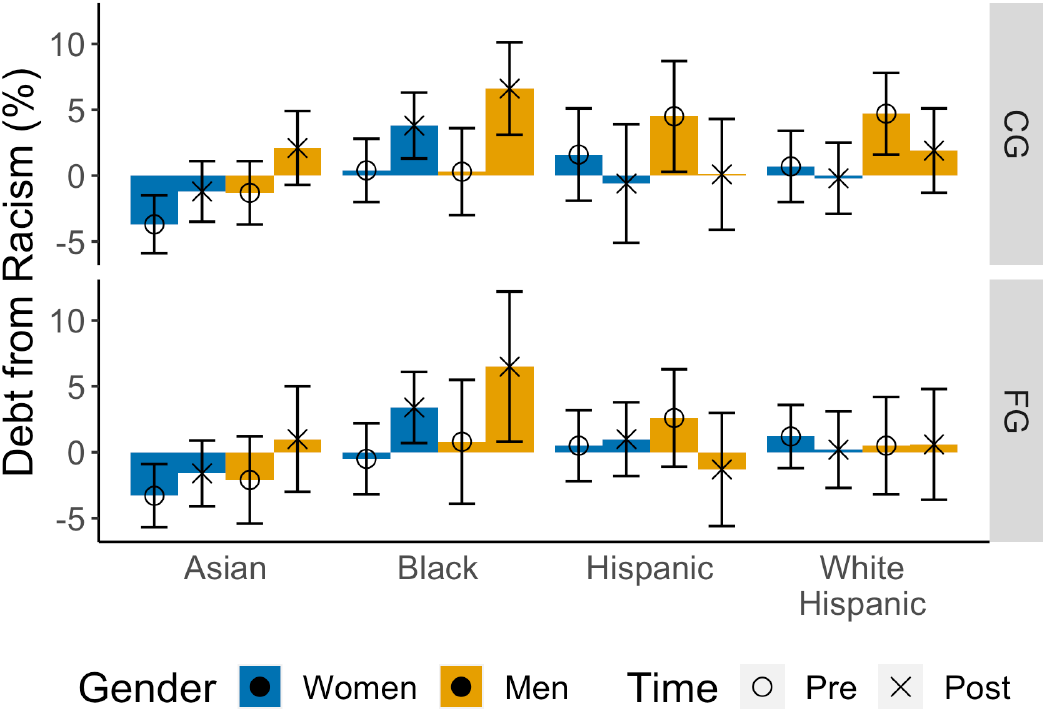
Comparing estimated scores for Asian, Black, Hispanic, and White Hispanic students to the White students with the same gender and first-generation identities. In this figure, positive values indicate larger societal educational debts. The error bars represent the addition of 1 standard error from each estimate. As discussed in the methods, we are very confident of the robustness of differences where the error bars do not include zero.

**Figure 7.**
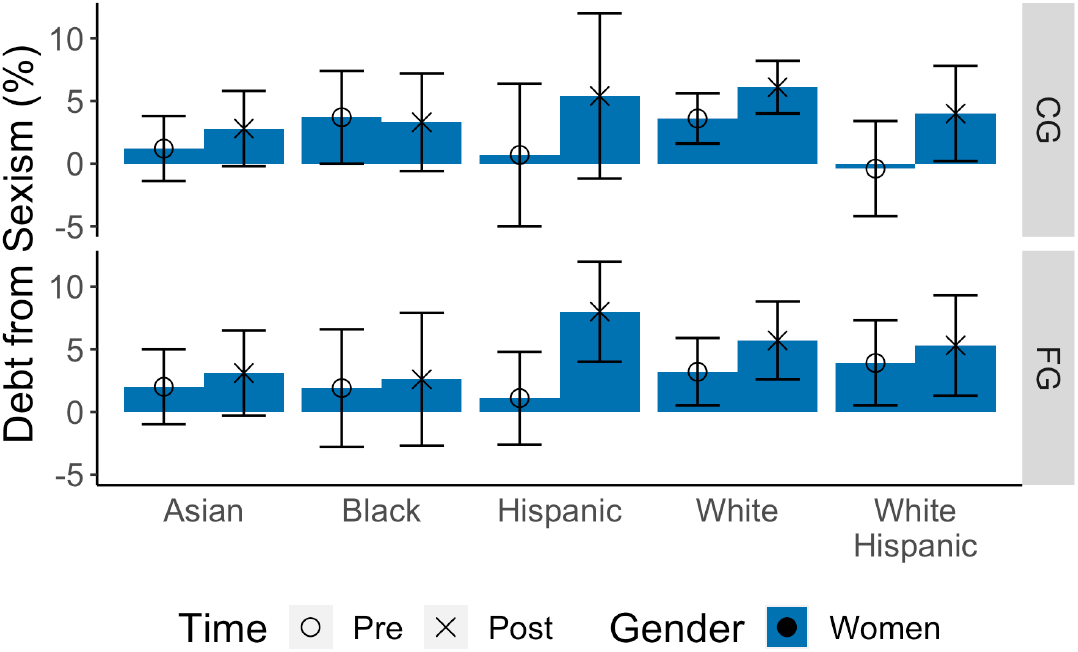
Comparing estimated scores for women and men with the same race and first-generation identities. In this figure, positive values indicate larger societal educational debts. The error bars represent the addition of 1 standard error from each estimate.

**Figure 8.**
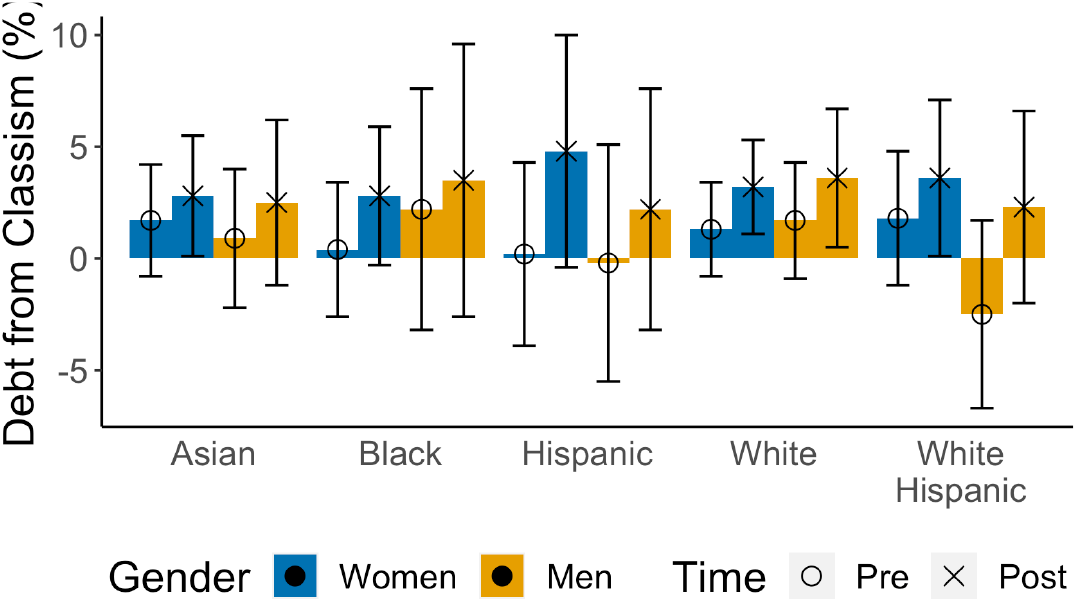
Comparing estimated scores for first-generation and continuing-generation students with the same gender and racial identities. In this figure, positive values indicate societal educational debts. The error bars represent the addition of 1 standard error from each estimate. These error bars are larger than those in Figures 6 and 7 because the comparisons almost always include a small group and only a subset of the students had data about their first-generation identity.

**Figure 9.**
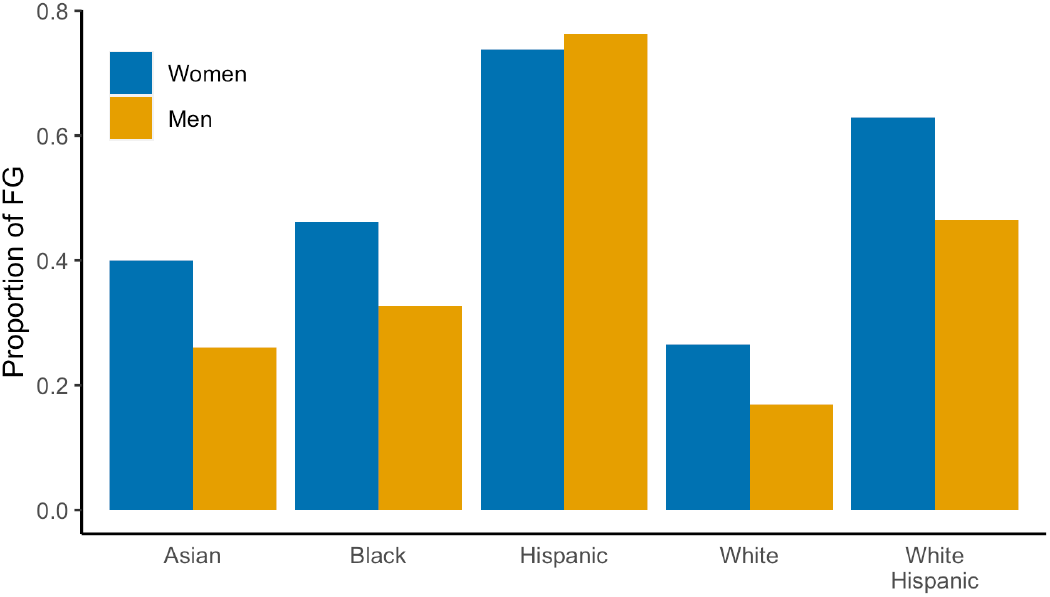
The proportion of first-generation college students by race and gender. The proportion varies across both race and gender.

**Figure 10.**
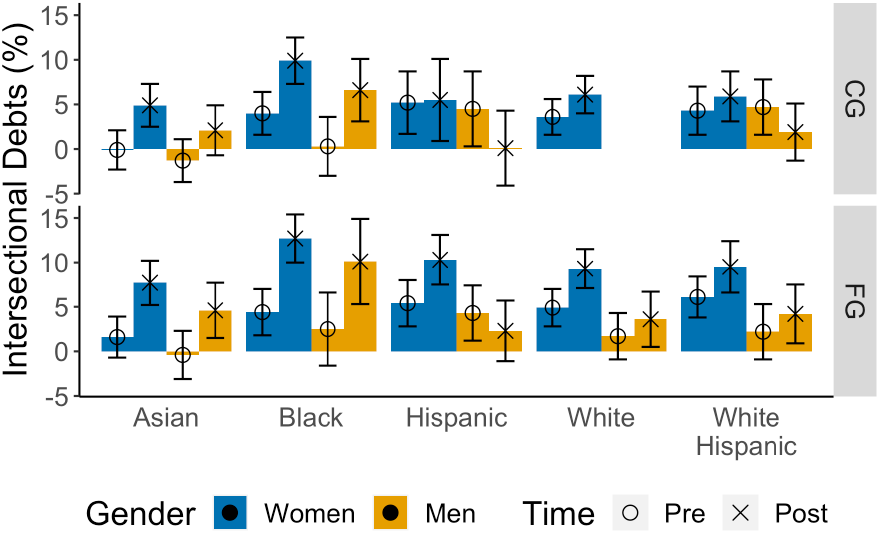
Comparing estimated scores for each group to continuing-generation White men. Positive values indicate society’s educational debts. Larger posttest educational debts than pretest for most groups indicate consistent increases in the educational debts society owes to students from marginalized groups. The error bars represent the addition of 1 standard error from each estimate.

## Findings

The findings section focuses on the predicted outcomes for each group (Table 2; Figure 5). We do not directly examine the model’s coefficients, which we include in the Supplemental Materials, because it requires combining up to 32 coefficients to predict a group’s score (e.g., First generation White Hispanic women’s posttest scores). To examine society’s educational debts, research questions 1 and 2, we first look at racism, sexism, and classism separately for the educational debts before and after instruction. These separate sections inform the consistency of the educational debts by only comparing across the one social identifier. We include separate figures for each of these comparisons of the differences and uncertainties between the two groups being compared. These figures came from the data in Table 3 but aid in making all the various comparisons. We then take an intersectional perspective, research question 3, that accounts for many social identifiers at once and how different identities tend to occur together.

### Society’s Educational Debts

#### Racism

We made sixteen comparisons, shown in Figure 6 on the pretest and posttest comparing scores for Asian, Black, Hispanic, and White Hispanic students to White students with the same gender and first-generation identities. On the pretest, 11 comparisons showed a societal educational debt owed to Hispanic students, White Hispanic students, and Black students except for first-generation Black women. The scores for all groups of Asian students and first-generation Black women were higher than for the similar group of White students. Most of these educational debts were relatively small and well within the uncertainty of the measurements. The educational debts for Hispanic and White Hispanic continuing-generation men were larger than the uncertainty (the combined standard errors for both measures), however, the total number of continuing-generation Hispanic men was only nine. These results do not indicate a large educational debt due to racism before instruction.

On the posttest, 11 of the 16 comparisons indicated a societal educational debt. Two differences stand out from the pretest data. First, the data indicated an educational debt owed to all four groups of Black students that was larger than the uncertainty in the measurements. These educational debts ranged between 3.4% and 6.6%. Second, the higher scores on the pretest for Asian students than White students all decreased to the posttest, and all differences were within the uncertainty of the measurement. Third, the differences for all groups other than Black students tended to be small. The largest showed an educational debt of 2.1% for continuing-generation Asian men and 1.9% for continuing-generation White-Hispanic men.

Making the same comparisons for the shifts in scores from pretest to posttest indicated similar results. The shifts indicated that the educational debts increased for Black (by 3.4% to 6.3%) students and the higher scores for Asian students on the pretest decreased (by 1.7% to 3.4%). The educational debts stayed the same or decreased for Hispanic and White-Hispanic students (from a 0.5% increase to a 4.4% decrease). These results indicated that instruction added to the societal educational debts owed to Black students, mitigated debts owed to Hispanic and White Hispanic students, and decreased the difference between Asian and White students.

#### Sexism

Comparisons, as shown in Figure 7, on the pretest showed educational debts owed to nine of the ten groups of women (between -0.4% and 3.9%). The posttest showed educational debts owed to all ten groups of women (between 2.6% and 8%). The educational debts increased for nine of the ten groups (from -0.4% to 6.9%). While only three comparisons were larger than the uncertainty on the pretest and five on the posttest, the results indicated a consistent educational debt owed to women and that instruction added to society’s educational debt.

#### Classism

For the pretest, as shown in Figure 8, eight of the ten comparisons showed educational debts owed to first-generation students (between -2.5% and 1.7%). None of these differences, however, were larger than the uncertainties in the measurements. Comparisons on the posttest showed educational debts for all ten comparisons (between 2.2% and 4.8%) with four of the comparisons larger than the uncertainty in the measurement. The educational debts increased for all ten groups (by 1.1% to 4.8%). The consistency of these results indicated that instruction increased the educational debt society owed to first-generation students.

#### Intersectionality

For the intersectional comparison, we compared each group to continuing-generation White men because the literature review indicated these were the most advantaged and least marginalized intersecting identities. We maintained this comparison even though continuing-generation White men had slightly lower scores than continuing-generation Asian men (1.3%) and women (0.1%) and first-generation Asian men (0.4%) on the pretest because they ended with the highest scores of any group on the posttest.

The descriptive data, shown in Figure 9, illustrates the need for an intersectional perspective that accounts for race, gender, and SES. Women were much more likely than men to identify as first-generation students. White students were much less likely than Asian, Black, Hispanic, and White Hispanic students to identify as first-generation students; Hispanic students were the most likely to identify as first-generation. These differences in the frequency of intersecting identities indicate that comparisons across race, gender, or SES alone may produce misleading findings.

The educational debts prior to instruction, shown in Fig. 10, ranged between -1.3% and 6.1% with a median of 4%. Sixteen of the nineteen educational debts increased from pre to post instruction. The three that decreased were for continuing- and first-generation Hispanic men and continuing-generation White Hispanic men. Society owed the largest educational debts after instruction to first-generation Black (12.7%) and Hispanic (10.3%) women. Society owed the next two largest educational debts to first-generation Black men (10.1%), the only group of men with such large educational debts, but a group with only 17 students total, and continuing generation Black women (9.9%), the only group of continuing generation students with such large educational debts. These results show the compounding impacts of anti-Black racism, sexism, and classism. After instruction, society owed the smallest educational debts to Hispanic, White-Hispanic, and Asian continuing-generation men. The small number of Hispanic continuing generation men, nine, means that these results are very uncertain for this group. Society owed the next smallest educational debts to Hispanic, White, White-Hispanic, and Asian first-generation men. The educational debts for these first-generation men were still quite large (between 2.3% and 4.6%). These results show that introductory biology courses greatly added to the educational debts owed to Black students and women, especially those that were also first-generation students.

## Limitations

While the LASSO platform that provided the data offers a large database that includes courses with more diverse scores than the literature (Nissen et al., 2020), it still underrepresents the institutions that disproportionately serve marginalized students (e.g., historically Black colleges and universities, Hispanic-serving institutions, and 2-year colleges). While we aren’t aware of any statistics on the breakdown of students in introductory biology courses by race, the data has a higher proportion of Black, Hispanic, and Asian students (9.7, 16.9, and 16.4%) than their proportion in the STEM degrees awarded nationally (8.6, 12.1, and 11.6%). The findings, therefore, may not represent instruction in two-year colleges but provide a reasonable proxy for STEM education at Bachelor’s degree-granting institutions with the following limitations.

Because the Learning Assistant Alliance hosts the LASSO platform (Otero et al., 2016), 54% of the courses used LAs (Otero, 2015; Barrasso & Spilios, 2021) to engage students in collaborative learning. The LASSO data also introduces potential selection bias, as instructors must know about LASSO and be interested in using a concept inventory to assess their course. This bias makes it likely that the instructors are more aware of research-based teaching practices and resources than the average instructor. Almost all the instructors (85 of 87) self-reported engaging students in collaborative learning in their courses. The findings do not speak to the status of lecture-based courses.

While the IMCA has undergone several rounds of validation research, it was developed at a highly competitive, primarily White institution and has not undergone analysis to test for race or gender bias (e.g., DIF analysis). Expansion of the validation argument to include diverse student groups would strengthen IMCA data’s ability to support claims about equity in student learning (Padilla, 2004).

The size of several of the groups for first- or continuing-generation were relatively small. The smallest group, continuing-generation Hispanic men, only included nine respondents. Small numbers create uncertainty about the societal educational debts owed to these groups. Additional research is necessary to confirm the results for continuing-generation Hispanic men in particular.

## Discussion

To contextualize the magnitude of the educational debts we found in our analyses, we follow the advice of Lortie-Forgues, Sio, and Inglis (2021) and discuss them in terms of the equivalent instruction time they represent. We use the overall average increase in scores from pretest to posttest of 16.3 percentage points as a proxy for the learning that occurs in a term. For example, an educational debt of 8.2 percentage points would be the equivalent of half of the learning that occurred during a term of instruction. We found that society owed the largest debt to first-generation Black women (12.7%); this debt is approximately 80% of the learning (16.3%) that occurred in the term. Before instruction, society owed first-generation Black women a noteworthy (4.4%) but much smaller educational debt compared to after instruction. Similar trends occurred for almost every group with the largest debts owed to those students with two or more marginalized identities: particularly continuing-generation Black women, first-generation Hispanic women, and first-generation Black men. Society owed these four groups educational debts ranging between 60% and 80% of the learning that occurred during the term.

The consistent trend of increased educational debts shows that these introductory biology courses using student-centered and learning assistant-supported instruction added to the educational debts society owed these students due to racism, sexism, and classism. Student-centered and learning assistant-supported instruction often provides the same opportunities and resources to all the students in a course: more collaboration, more instructor and learning assistant contact, more guidance and feedback, and less lecturing. These results indicate that students with multiple marginalized identities benefit less from these instructional practices in biology than students with few or no marginalized identities in terms of changes in conceptual knowledge. As detailed in our definition, these are racist, sexist, and classist outcomes because they add to the power differences between continuing-generation White men and other groups. This contrasts prior findings of learning assistant-supported and student-centered instruction in chemistry repaying society’s educational debts (Van Dusen, et al., 2021) and similar physics instruction increasing conceptual learning compared to lecture based instruction but still maintaining educational debts (Van Dusen & Nissen, 2020). Providing the same resources to everyone will only repay society’s educational debts if the students owed those debts benefit more from the resources than White students, continuing-generation students, and men.

Researchers have proposed several solutions to mitigate the impacts of racial, gender, and class disparities on marginalized students. Several studies indicate that active learning, specifically high-intensity active learning (>66% class time spent on active learning) may repay educational debts as measured by exam performance (42% reduction) and course passing rates (76% reduction) owed to Black, Hispanic, and low-SES students across the STEM disciplines (Theobald et al., 2020). While active learning may generally help lower educational debts compared to traditional lecture instruction, our findings show an increase in debts post-instruction. The difference between Theobald et al. and our findings may come from several factors. Theobald et al’s data restricted them to analyze URM students and low-SES students as one group and the analysis did not include gender. The data in Theobald et al. included a mixture of test types, aggregated data across disciplines, and included upper and lower division courses. These choices allowed Theobald et al. to answer the important question about how much active learning is necessary to support equity. In contrast, our study only looked at one concept inventory in introductory biology courses looking at the intersection of racism, sexism, and classism in data from courses that almost all used collaborative instruction (85/87) and half used learning assistants (47/87). It is possible that active learning could reduce the differences between groups compared to lecture-based instruction while still adding to the educational debts due to racism, sexism, and classism that students start courses with. This result would mean that active learning adds to these debts less than lecture-based instruction. This improvement represents a valuable first step, but only a first step. As Theobald et al. (2020) state, “…active learning is not a silver bullet for mitigating achievement gaps” (p. 6479). Nor is it a silver bullet for repaying the educational debts society owes to Black, Hispanic, White Hispanic, women, and first-generation students.

Our figures and the common language of achievement gaps used in the prior quote give the false impression of chasms separating the scores of different groups. The results in our descriptive statistics, Supplemental Figure 1, show that these differences are not gaps that separate groups but differences in the typical scores of students within each group. Many Asian, Black, Hispanic, and White Hispanic men and women and White women had perfect scores on the assessment. The societal educational debts lie in the distributions of scores, not in a gap that separates one group from another.

Society’s educational debts, accrued over the course of a student’s undergraduate tenure, likely contribute to the greater attrition of students from marginalized groups. Results show that Pell grant recipients, an indicator of lower SES associated with material capital, were more likely to leave STEM majors at the Bachelor’s level than non-recipients (Chen, 2013). We could not find studies speaking specifically to marginalized groups in undergraduate biology majors. While research should investigate attrition across STEM majors and intersectional student identities, we find it reasonable to infer that the addition of educational debts leads to greater attrition amongst women, first-generation students, and Black and Hispanic students. This attrition can feed into a cycle of talent loss and lower representation in the field, which adds additional challenges to students that decide to stay in the biology major because they don’t see many people like them succeed and join the biology workforce.

The intersectional educational debts that society owes students with overlapping marginalized identities found in this study point to the need for intersectional research. Because oppressive ideologies intertwine, research cannot completely disentangle them. Researchers, however, should account for how different oppressive structures emanating from different ideologies may have disproportionate effects on students. We treat first-generation status as a marker of SES because students with any gender and any race can be first-generation students. The data showed that the proportion of first-generation students varied across race and gender. Seventy-four percent of Hispanic women and men reported being first-generation college students, the highest proportion of any group, followed by White Hispanic students. These results indicate that classism may play a role in the educational debts owed to Hispanic and White Hispanic students distinct from racism. Interventions and structural transformations focused on the impacts of racism separately from classism may not repay the educational debts owed to Hispanic and White Hispanic students. Discipline based education research seldom looks at class or SES (Talbot, 2021). When it does, discipline based education research on SES seldom defines SES or establishes its work within a theoretical perspective on SES (Talbot, 2021). These practices leave few findings for us to draw on that are specific to classism in college STEM instruction, classism in college biology instruction, or the intersections of racism, classism, and sexism in biology instruction.

## Conclusion

The broad systemic nature of racism, sexism, and classism makes it difficult for them to inform interventions or overhauls of the educational systems that create inequitable outcomes. We use them, rather, to make explicit that the inequalities we measured in this data represent the outcomes of oppressive ideologies and structures. To structure our conclusion, we draw on the four I’s of oppression discussed in the Conceptual Framework. The implicit and explicit *ideologies* that communicate messages about who belongs in biology and where they belong in the field. And the three ways in which these ideologies manifest: *institutional, interpersonal*, and *internalized oppression* (Paradies, 2006).

Research often esteems the biological sciences as a field that supports gender equity (Cheryan et al., 2017). Our results, however, indicated that introductory university biology courses added to the educational debts society owed to women, Black, Hispanic, White Hispanic, and first-generation students due to the intersection of racism, sexism, and classism. Society owed the largest debts after instruction to students with multiple marginalized identities: Black and Hispanic, first-generation women. These results align with the national statistics showing that women and Black and Hispanic Americans make up a much smaller proportion of people working or earning a PhD in the biological sciences than those earning a bachelor’s degree in the biological sciences.

We used racism, classism, and sexism and societal educational debts in this paper to refute the idea that these differences represent deficits of the students. The trends in populations we describe here result from a long history of systemic oppression ingrained in society. The responsibility to address these systems lies most heavily on those with the greatest power to make change; administrators, educators, and education researchers carry much greater power to make changes than the undergraduate students whom our study focused on. The increasing educational debts we found indicate a need for individual and institutional action to address these inequities. Institutional, interpersonal, and internalized oppression can frame research on and action in biology education to repay rather than add to the educational debts society owes students from marginalized groups.

Ideologies and institutional, interpersonal, and internalized oppression are not independent. They interact and reinforce one another. They can, however, provide a framework for anti-oppressive work that takes multiple approaches. Research can approach investigating both the implicit and explicit ideologies (Gawronski, 2018) held by instructors, administrators, scientists and students in disciplines within the biological sciences to identify harmful ideologies, such as the association of brilliance with men (Leslie et al., 2015) or that students have fixed abilities (Canning et al., 2019). Oppressive ideologies may not occur across all the biological sciences but may occur often within some disciplines. At the institutional level, introductory science courses often fail to support students who could succeed but have had fewer or poorer prior opportunities in the sciences (Seymour and Hewitt, 1997; Seymour and Hunter, 2019). Biology courses and degree programs could instead use inspiration from universal design (Scanlon et al., 2018; Edyburn, 2010) to develop courses and programs that support students with diverse interests, backgrounds, and identities to pursue and succeed in careers in the biological sciences. Educators can adopt reflective practices from curriculum designed to counter racism and sexism, such as the Underrep Curriculum (Underrep project, 2022; Doucette et al., 2021), to assist their students in reflecting on who gets to do biology and how that has or has not changed over time to take on the ideologies at the root of these results.

Future research should investigate how institutionalized racism and classism in funding for primary and secondary education lays the foundation for these educational debts that may be added to by policies in higher education institutions around remedial math and English courses, college entrance exams, and introductory STEM courses that fail the students who come with fewer prior opportunities instead of preparing them for success. At the level of personal interactions, further work should focus on the kinds of interactions that students have with each other, classroom assistants, and instructors in these collaborative courses that may add to these educational debts: from ignoring students of color to microaggressions and outright aggression. Work to address internalized oppression can focus on ways in which the learning environment disproportionately harms oppressed groups. Exploring how courses can support students in developing identities as biologists and scholars, in seeing their hard work support their success (e.g., growth mindsets), and creating environments that do not trigger stereotype threats will ultimately help develop strategies to repay society’s educational debts.

While this work does not have sufficient lecture-based courses to show that collaborative instruction is or is not more equitable than lecture-based courses, several studies show that students learn more and get better grades in collaborative, student-centered courses (Theobald et al, 2020; Freeman et al., 2014; Hake, 1998; Van Dusen and Nissen, 2020). While our results show that collaborative introductory biology courses tend to add to society’s educational debts, that does not mean they can’t or don’t in some cases repay these educational debts. These collaborative courses will likely provide more useful insights for how to best repay the educational debts that society owes due to racism, sexism, and classism than lecture-based courses.

Data from 87 introductory biology courses at 11 institutions that primarily used collaborative instruction, often with the support of learning assistants, indicated that these courses added to the educational debts society owed students with marginalized identities. Society owed the largest educational debts after instruction to students with multiple marginalized identities equal to four-fifths of the learning that occurred during a semester. These results, combined with how SES varies across racial groups and genders, show that intersectional analyses and the datasets that enable them provide necessary information for building and maintaining equitable biology instruction.

## Acknowledgments

This work was funded in part by NSF-IUSE Grants No. DUE-1928596.

## Footnotes

1 In this publication, we capitalize all races, including White, emphasizing that there is no default race and that they are all social constructs with associated sets of cultural practices.

2 Racism, sexism, and classism are not the only forms of oppression and our focus on them alone results from the limitations of this study.

## Notes

### Competing Interest Statement

The first and second authors are the directors of the LASSO platform, which was used for the data collection.

