## Supplemental Material for "A QuantCrit investigation of society’s educational debts due to racism, sexism, and classism in biology student learning"

This document is comprised of two sections:
1. Supplemental Tables and Figures
2. Demographic Survey.

**Section 1: Supplemental Tables and Figures**

**Supplemental Table 1.** Descriptive statistics for the data overall and by Race and Gender. Descriptive statistics for the subset of students who responded to the question about their first-generation status is provided below. The Ns in each column do not add to the total. We do not include descriptive statistics for the other gender and other race groups and students who identified as more than one race are included in the descriptive statistics for each of those races.

| | | | N | | Mean | | Std. Dev. | |
|---|---|---|---|---|---|---|---|---|
| Race | Gender | Total | Pre | Post | Pre | Post | Pre | Post |
| All | All | 6547 | 5803 | 4387 | 44.0 | 60.3 | 16.8 | 18.2 |
| AIAN | Man | 32 | 26 | 20 | 41.2 | 65.2 | 13.8 | 17.2 |
| AIAN | Woman | 62 | 53 | 38 | 41.4 | 62.2 | 14.6 | 20.3 |
| Asian | Man | 355 | 317 | 224 | 48.3 | 63.8 | 17.4 | 17.2 |
| Asian | Woman | 717 | 651 | 533 | 46.6 | 61.6 | 17.0 | 17.8 |
| Black | Man | 141 | 119 | 82 | 44.8 | 58.2 | 18.9 | 19.2 |
| Black | Woman | 494 | 424 | 327 | 43.4 | 56.4 | 16.3 | 17.4 |
| Hispanic | Man | 132 | 113 | 81 | 39.3 | 61.9 | 16.0 | 16.6 |
| Hispanic | Woman | 270 | 236 | 161 | 39.4 | 54.4 | 14.8 | 20.0 |
| PacIsland | Man | 20 | 17 | 11 | 41.7 | 64.0 | 12.7 | 18.3 |
| PacIsland | Woman | 46 | 38 | 33 | 41.8 | 63.0 | 16.7 | 19.5 |
| White | Man | 1160 | 1008 | 741 | 46.3 | 65.4 | 18.1 | 17.9 |
| White | Woman | 2496 | 2236 | 1761 | 42.6 | 59.0 | 16.0 | 18.1 |
| White Hispanic | Man | 189 | 163 | 109 | 43.5 | 62.9 | 18.1 | 17.8 |
| White Hispanic | Woman | 488 | 447 | 309 | 40.5 | 57.4 | 15.7 | 17.7 |

**Supplemental Figure 1.** Violin plots, boxplots, and scatter plots of the data for pretest and posttest IMCA scores. The violin plot is a reflected density plot to show the distribution of the data. The notched boxplots show the distribution but focus attention on the medians with notches to show the 95% confidence intervals. The scatter plot is jittered to randomly distribute the points left and right of the vertical axis for clarity. The scatter plot illustrates the number of data points in each group and adds detail to how the data is distributed, particularly in the tails. These plots show easily identifiable differences across groups, but also show these differences are shifts in the distributions of scores and not gaps that separate groups.

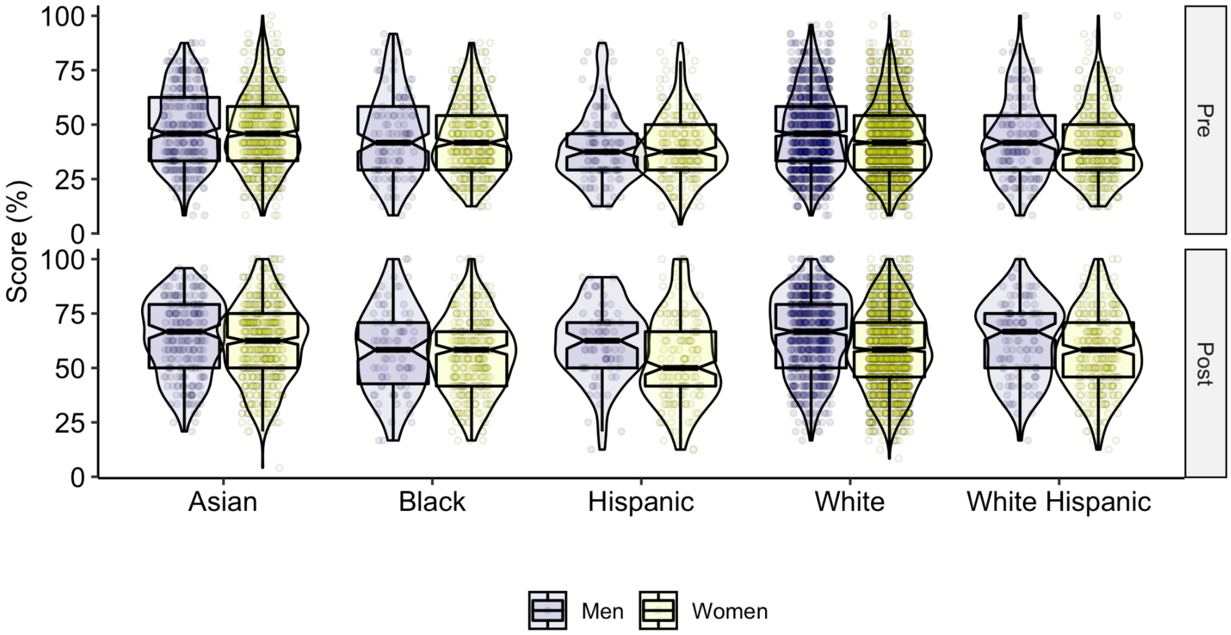

**Supplemental Table 2.** Descriptive statistics by race, gender, and first-generation status for the subset of students with data for first-generation status.

| | | First | | N | | Mean | | Std. Dev. | |
|---|---|---|---|---|---|---|---|---|---|
| Race | Gender | Generation | Total | Pre | Post | Pre | Post | Pre | Post |
| All | All | Yes | 955 | 813 | 582 | 40.3 | 58.8 | 15.9 | 20.0 |
| All | All | No | 1867 | 1603 | 1298 | 42.7 | 62.5 | 16.6 | 18.7 |
| Asian | Man | Yes | 30 | 25 | 21 | 46.2 | 62.3 | 18.3 | 19.2 |
| Asian | Man | No | 85 | 71 | 55 | 45.5 | 64.8 | 18.6 | 17.7 |
| Asian | Woman | Yes | 84 | 67 | 69 | 40.4 | 62.1 | 16.1 | 22.0 |
| Asian | Woman | No | 126 | 113 | 96 | 44.0 | 65.1 | 17.6 | 17.7 |
| Black | Man | Yes | 17 | 14 | 9 | 38.4 | 48.1 | 11.5 | 21.3 |
| Black | Man | No | 35 | 32 | 18 | 40.1 | 51.6 | 17.5 | 19.8 |
| Black | Woman | Yes | 91 | 76 | 49 | 40.6 | 57.9 | 14.5 | 21.2 |
| Black | Woman | No | 106 | 87 | 68 | 41.5 | 59.4 | 17.3 | 18.6 |
| Hispanic | Man | Yes | 29 | 28 | 14 | 39.0 | 64.0 | 13.3 | 20.2 |
| Hispanic | Man | No | 9 | 6 | 5 | 32.6 | 58.3 | 12.8 | 7.8 |
| Hispanic | Woman | Yes | 73 | 64 | 38 | 37.9 | 60.9 | 11.7 | 23.0 |
| Hispanic | Woman | No | 26 | 24 | 13 | 37.8 | 66.7 | 16.3 | 21.9 |
| White | Man | Yes | 89 | 79 | 55 | 41.0 | 62.1 | 17.7 | 20.4 |
| White | Man | No | 437 | 355 | 305 | 45.3 | 67.5 | 17.8 | 18.3 |
| White | Woman | Yes | 299 | 246 | 202 | 40.5 | 56.1 | 15.9 | 18.9 |
| White | Woman | No | 827 | 726 | 599 | 41.8 | 60.2 | 15.6 | 18.6 |
| White Hispanic | Man | Yes | 46 | 38 | 23 | 42.1 | 62.5 | 20.4 | 19.6 |
| White Hispanic | Man | No | 53 | 44 | 30 | 38.3 | 62.8 | 13.9 | 17.7 |
| White Hispanic | Woman | Yes | 161 | 145 | 84 | 38.5 | 58.2 | 15.5 | 19.5 |
| White Hispanic | Woman | No | 95 | 89 | 61 | 41.0 | 61.9 | 16.5 | 17.7 |

**Supplemental Table 3.** The AICc scores and variables removed from each of the ten best models. Model 1 had the lowest AICc score and was used in our analysis. Model 2 also fit the data well, but all of the other models were at more than 2 AICc points higher than Model 1. In these models, the "test" variable connotes the posttest.

| Model | AICc | Delta AICc | Variables removed | | | |
|---|---|---|---|---|---|---|
| | | | retake*test | Black*FG* Test | Black*test* women | Black*FG*test* women |
| 1 | 110823.7 | 0.0 | | | | |
| 2 | 110824.0 | 0.3 | x | | | |
| 3 | 110827.5 | 3.8 | | | x | |
| 4 | 110827.8 | 4.1 | x | | x | |
| 5 | 110829.8 | 6.1 | | x | x | |
| 6 | 110830.1 | 6.4 | x | x | x | |
| 7 | 110830.2 | 6.5 | | | x | x |
| 8 | 110830.5 | 6.8 | x | | x | x |
| 9 | 110832.4 | 8.7 | | x | x | x |
| 10 | 110832.7 | 9.0 | x | x | x | x |

**Supplemental Table 4.** The model coefficient and errors for the final model.

| Coefficient | Est. | Est. Error | | Coefficient | Est. | Est. Error |
|---|---|---|---|---|---|---|
| Intercept | 46.97 | 1.43 | | women*Black | -2.50 | 2.51 |
| retake | 0.72 | 0.73 | | women*Hispanic | 0.53 | 4.25 |
| test | 16.34 | 1.66 | | women*White | -2.38 | 1.59 |
| gender_other | 4.75 | 2.51 | | test*gender_other | -2.89 | 3.51 |
| race_other | -2.00 | 1.17 | | test*race_other | -0.79 | 1.47 |
| FG | -0.85 | 2.02 | | test*FG | -1.56 | 2.65 |
| women | -1.15 | 1.40 | | test*women | -1.68 | 1.84 |
| Black | -1.60 | 2.26 | | test*Black | -2.75 | 3.44 |
| Hispanic | -5.80 | 3.34 | | test*Hispanic | 7.82 | 4.16 |
| White | -1.30 | 1.29 | | test*White | 3.40 | 1.77 |
| FG*women | -0.89 | 2.53 | | FG*Hispanic*White | 3.19 | 5.98 |
| Hispanic*White | 1.13 | 4.02 | | women*Hispanic*White | 3.46 | 5.17 |
| FG*Black | -1.39 | 4.31 | | FG*women*Black | 2.77 | 4.61 |
| FG*Hispanic | 1.04 | 4.45 | | FG*women*Hispanic | 0.46 | 5.82 |
| FG*White | -0.84 | 2.73 | | FG*women*White | 1.25 | 3.38 |
| test*FG*women | 0.53 | 3.01 | | test*women*White | -0.89 | 2.00 |
| test*Hispanic*White | -5.10 | 5.13 | | FG*women*Hispanic*White | -5.28 | 7.65 |
| test*FG*Black | 0.02 | 5.78 | | test*FG*Hispanic*White | -2.09 | 7.46 |
| test*FG*Hispanic | -0.88 | 5.11 | | test*women*Hispanic*White | 1.15 | 6.08 |
| test*FG*White | -0.28 | 3.05 | | test*FG*women*Black | -1.46 | 6.20 |
| test*women*Black | 1.88 | 3.78 | | test*FG*women*Hispanic | -2.82 | 6.16 |
| test*women*Hispanic | -3.01 | 5.36 | | test*FG*women*White | -0.62 | 3.58 |
| | | | | test*FG*women*Hispanic*White | 6.04 | 8.47 |

**Section 2: Demographics Survey**

The demographics questions and options asked by the LASSO platform in the Spring of 2022.

How do you describe your gender? (check all that apply)
Options: Man, woman, transgender, genderqueer/gender non-conforming, another identity (please state), and prefer not to answer.

What is your ethnicity?
Options: Not of Hispanic/Latinx/Spanish origin, Mexican/Mexican American/Chicano/Chicana, Puerto Rican, Cuban, another Hispanic/Latinx/Spanish origin (please state), and prefer not to answer.

What is your race? (check all that apply)
Options: White, Black or African American, Chinese, Filipino, Asian Indian, Vietnamese, Japanese, Korean, an Asian race not listed, Native Hawaiian, Samoan, Chamorro, a Pacific Islander race not listed, Middle Eastern or North African, another race/other races (please state), and prefer not to answer.

Have you been a Learning Assistant in another course?
Options: Yes and No.

What is your role in the class?
Options: student, learning assistant, teaching assistant, and faculty.

What is your status in school?
Options: 1st year, 2nd year, 3rd year, 4th year, 5th year, 6th year, and graduate student.

Is this your first time taking the course?
Options: yes and no.

Have any of your parents or guardians received a degree from a 4-year college?
Options: yes, no, I do not know, and prefer not to answer.
